# Improved recombinant protein production and scale-up fermentation of *Aspergillus oryzae* hyphal-dispersion hydrophobin-deficient strain

**DOI:** 10.64898/2026.08.06.739278

**Authors:** Shunya Susukida, Yasuhiro Baba, Makoto Fujisawa, Yuki Niikawa, Kiyoaki Muto, Ken Miyazawa, Akira Yoshimi, Yoshikazu Kato, Hirofumi Horiguchi, Keietsu Abe

## Abstract

In liquid fermentation of filamentous fungi such as *Aspergillus oryzae*, increased broth viscosity and biomass adhesion to bioreactor surfaces remain major challenges. We previously developed a hyphal dispersion mutant lacking two hyphal adhesion factors, namely cell wall α-1,3-glucan (AG) and biofilm galactosaminogalactan (GAG) (AGΔ-GAGΔ strain). The culture broth of the AGΔ-GAGΔ strain has low viscosity, which improves mixing and enzyme production. However, mycelia still extensively attach to bioreactor walls and downstream equipment, which impairs mixing and reduces product recovery. The hydrophobin RolA, a surface-active protein of *A*. *oryzae*, densely coats conidia and hyphae and contributes to cell surface hydrophobicity. In this study, we disrupted the *rolA* gene in AGΔ-GAGΔ (AGΔ-GAGΔ-Δ*rolA* strain) and evaluated the effects of this disruption on hyphal adhesion to the walls of culture vessels, enzyme production, and bioreactor performance. At the flask scale, the adhesion to glass surfaces was significantly reduced and recombinant enzyme activity was increased by 10%. Improved culture recovery at the end of fermentation further increased total enzyme yield. In a lab-scale stirred-tank bioreactor, both growth and enzyme production were increased. Scaling-up to a 200-L bioreactor showed reduced agitation power consumption while improving hydrodynamic properties. Fermentation of AGΔ-GAGΔ-Δ*rolA* was successfully scaled up to a 3000-L bioreactor; consistent enzyme activity and improved flow circulation in the bioreactors were confirmed by computational fluid dynamics analysis. Overall, the AGΔ-GAGΔ-Δ*rolA* strain has increased enzyme production and scalability, supporting its suitability for industrial applications.

## 1 Introduction

Industrial liquid fermentation of filamentous fungi faces several challenges that limit productivity such as increased broth viscosity due to filamentous morphology, difficulty in controlling macro-morphology, and biomass attachment to bioreactor surfaces and downstream equipment (Noorman, 2011). This attachment, referred to as wall growth, results in dense, crust-like mycelial layers that are often caused by a broth’s pseudoplasticity and foaming buildup, and impairs mixing and reduces effective working volume (Wongsamuth and Doran, 1994). Wall growth often contains necrotic cells, which can detach into the broth, lowering the fraction of active biomass and overall process efficiency (Piehl et al., 1988). Clogging of pipes and ports and unintended sporulation can further complicate operation, particularly when handling genetically modified strains, where preventing environmental dissemination and maintaining physical containment are important considerations (Kimman et al., 2008).

Hydrophobins are small amphipathic proteins produced by fungi. Class I hydrophobins, including *Aspergillus oryzae* RolA, self-assemble into amyloid-like fibrils known as rodlets that form hydrophobic surface layers (Wosten et al., 1993; Hektor and Scholtmeijer, 2005; Terauchi et al., 2020). These structures contribute to interfacial interactions, including adhesion to solid surfaces and are stabilized at hydrophilic–hydrophobic interfaces (Wosten et al., 1994; van Wetter et al., 2000).

We previously identified two polysaccharides, cell wall α-1,3-glucan (AG) and galactosaminogalactan (GAG) in the extracellular matrix, as hyphal adhesive factors and disrupted the genes required for AG and GAG biosynthesis in *A. oryzae*. In comparison with the wild-type strain, the resulting hyphal-dispersion strain (AGΔ-GAGΔ) had reduced broth viscosity and increased enzyme productivity because of the improved hydrodynamics (Miyazawa et al., 2019; Ichikawa et al., 2022; Susukida et al., 2025). However, this AGΔ-GAGΔ strain still had considerable wall growth under high-cell-density conditions.

In this study, we hypothesized that disruption of the *rolA* gene in this background would reduce biomass adhesion to the surface of culture vessels and improve fermentation performance. We evaluated the resulting strain in shake flasks and lab- and pilot-scale bioreactors by examining enzyme productivity, rheology, and mixing behavior, supported by computational fluid dynamics (CFD) analysis. The strain was successfully scaled up to 3000 L to assess its industrial applicability.

## 2 Materials and Methods

### 2.1 Medium

For inoculum preparation, Czapek-Dox agar plates and malt agar plates used as described previously (Miyazawa et al., 2016) except 2% KCl was added to malt agar medium.

For flask fermentation, a modified version of chemically defined medium (Diano et al., 2009) was used. The medium contained 3% glucose, 1% K_2_HPO_4_ buffer (pH 7.8), 0.3% NaNO_3_, 0.035% (NH_4_)_2_SO_4_, 0.1% MgSO_4_·7H_2_O, 0.1% NaCl, and 0.1% Tween 80. Trace elements were added to final concentrations of 7.2 mg L^−1^ ZnSO_4_‧7H_2_O, 1.3 mg L^−1^ CuSO_4_‧5H_2_O, 0.3 mg L^−1^ NiCl_2_‧6H_2_O, 3.5 mg L^−1^ MnCl_2_, and 6.9 mg L^−1^ FeSO_4_‧7H_2_O.

For lab-scale bioreactor fermentation, the medium consisted of 3% glucose, 1% peptone, 0.5% yeast extract, 0.5% meat extract, 20 mM succinate buffer (adjusted to pH 7.0 with NaOH), 0.5% K_2_HPO_4_, and 0.1% MgSO_4_·7H_2_O.

For pilot-scale bioreactor fermentation, the medium contained 2% glucose, 1.5% yeast extract, 1% K_2_HPO_4_, 0.3% NH_4_Cl, and 0.05% MgSO_4_·7H_2_O; pH was adjusted to 6.0 with diluted H_2_SO_4_.

For all fermentations, media were supplemented with 0.05% (v/v) emulsified silicone antifoam agent, 0.05% (v/v) non-ionic polyether antifoam agent, and 0.005% (w/v) chloramphenicol. Glucose, succinate buffer, MgSO_4_·7H_2_O, and chloramphenicol were sterilized separately and added after autoclaving for flask and lab-scale bioreactor cultures.

### 2.2 Strains

*Aspergillus oryzae* NS4 (*sC*^-^, *niaD*^-^) carrying Δ*ligD* (Δ*ligD*::*sC*, Δ*adeA*::*ptrA*) (Mizutani et al., 2008) and triple-deletion strain of three α-1,3-glucan synthases (AGΔ) (Miyazawa et al., 2016) were used for all genetic manipulations.

Disruption of the *rolA* gene was performed using the adenine auxotroph marker gene *adeA* following the methods described in Miyazawa et al. (2019). Briefly, to construct the WT-Δ*rolA* strain, a disruption cassette was generated using the NS4 strain. To generate the AGΔ-GAGΔ background, a Cre–*loxP* marker recycling system (Zhang et al., 2017) was applied to the AGΔ strain to introduce *adeA* and disrupt two genes in the galactosaminogalactan biosynthesis cluster (*sphZ* and *ugeZ*). Transformants were cultivated on xylose-containing agar to induce Cre recombinase, resulting in the AGΔ-GAGΔ strain with adenine auxotrophy. To construct the AGΔ-GAGΔ-Δ*rolA* strain, the same *rolA* disruption cassette was introduced.

Xylanase was chosen as a model recombinant enzyme (Ishibashi et al., 2026). WT-*xynF1* and AGΔ-GAGΔ-*xynF1* strains were used as reported by Ishibashi et al. (2026), and XynF1-overexpressing *rolA*-deficient strains were constructed accordingly. Hereafter, the four xylanase production strains are referred to as WT, WT-Δ*rolA*, AGΔ-GAG-Δ*rolA*, AGΔ-GAGΔ-Δ*rolA* (Table S1).

The RolA-overexpressing *A. oryzae* strain (Takahashi et al., 2005) was used to purify RolA as an antigen for polyclonal anti-RolA rabbit antibody production.

### 2.3 Fermentation

For flask fermentation, 500-mL Sakaguchi flasks (Sansho, Tokyo, Japan) containing 100 mL of medium were used. The final inoculum concentration was 5 × 10^5^ conidia/mL. Cultures were incubated at 30°C for 72 h on a shaker (BR-3000, Taitec, Saitama, Japan), with rotational agitation for 0–12 h followed by reciprocating agitation for 12–72 h, both at 125 rpm. Samples (10 mL) were collected at 48 and 72 h, and 10 mL of feeding medium (25% glucose, 4% urea) was added immediately after each sampling. To suppress excessive wall growth, attached biomass was scraped from the flask inner wall and suspended into the culture under sterile conditions after sampling at 48 h.

Lab-scale bioreactor fermentation was performed as described previously with a minor modification. A 4-L bioreactor, equipped with HS100 and HR impellers (d = 85 mm; Satake Multimix Corporation, Saitama, Japan), containing 2.1 L of medium was operated at 300 rpm for 0–24 h and 786 rpm thereafter, with an aeration rate of 1.68 L min^−1^. Cultures were maintained at 30°C for up to 120 h. The feeding medium (30% glucose, 0.5% peptone, 0.5% meat extract) was continuously supplied at 4 mL L^−1^ h^−1^ from 24 to 120 h. Aqueous NH_4_ (10%) was used to maintain pH at 5.0. Dissolved oxygen (DO) concentration, oxygen uptake rate (*OUR*), and volumetric power consumption under aeration (*P_gV_*) were determined as described previously (Susukida et al., 2025).

Pilot-scale fermentation of the AGΔ-GAGΔ-Δ*rolA* strain was conducted in 200-L and 3000-L bioreactors. At the 200-L scale, a single-use bioreactor (HSF-HSUB 200, Satake Multimix Corporation) equipped with HS100 and HR100 impellers (d = 265 mm) was used, with an initial working volume of 120 L. The inoculum (40 mL) was added to a final concentration of 5 × 10^4^ conidia/mL. The culture was agitated at 250 rpm for 0–24 h and 390 rpm thereafter, and aerated with ambient air at 96 L min^−1^.

At the 3000-L scale, fermentation was conducted at Green Earth Institute (Chiba, Japan). The bioreactor was equipped with one HS100 and two HR100 impellers (d = 540 mm) in a three-stage configuration. The initial working volume was 1800 L, and the inoculum (84 mL) was added to a final concentration of 5 × 10^4^ conidia/mL. Aeration speed was set at 1440 L min^−1^, and agitation was 150 rpm for 0–24 h and 263 rpm thereafter.

For both pilot scales, cultures were maintained at 30°C for 160 h. Feeding medium (50% glucose, 1.5% yeast extract) was supplied at 2.5 mL L^−1^ h^−1^ from 24 h to 144 h. The pH was maintained at 5.0 with 10% aqueous NH_4_ (200 L) or gaseous NH_4_ (3000 L). Agitation speed during 24–160 h was determined to achieve a *P_gV_* of approximately 3 kW m^−3^ during the final fermentation phase, as indicated by the CFD analysis described in 2.7.

### 2.4 Quantification of mycelial dry weight, enzyme activity, wall growth, and working volume

Culture samples (20 mL for the lab scale and 40 mL for the pilot scales) were collected. Aliquots (5 mL) were filtered through pre-weighed Miracloth (Merck Millipore, Darmstadt, Germany). The retained mycelia were washed twice with deionized water, dried, and weighed. XynF1 activity in the filtrate was measured as described by Ishibashi et al. (2026).

At the end of the fermentation in flasks and lab-scale bioreactors, the culture liquid was carefully aspirated using syringes or a vacuum pump without disturbing the wall growth. The recovered liquid volume was measured and defined as the final working volume, with correction for volume changes due to sampling and supply of feeding medium. To assess overall productivity, total enzyme activity and total biomass were calculated by multiplying the respective concentrations by the working volume. Wall growth was then carefully collected, filtered through pre-weighted Miracloth, washed, dried and weighed.

### 2.5 Immuno-fluorescent labelling of cell surface RolA

At the end of lab-scale fermentation, a 100-μL culture aliquot was gently mixed with 1 mL of phosphate buffered-saline (PBS), centrifuged at 15,000 × *g* for 1 min at room temperature, and decanted. The pelleted mycelia were washed once with 1 mL PBS. Polyclonal anti-RolA rabbit antibody produced by TK Craft (Nagano, Japan) as described by Takahashi et al. (2005) (1:200 dilution with PBS, 100 μL) was added to the pellet, gently mixed, and incubated at room temperature for 90 min with intermittent mixing. The samples were then washed three times with PBS and decanted. Subsequently, 50 μL of goat anti-rabbit IgG conjugated with Alexa Fluor 568 (1:5 dilution; Invitrogen, Carlsbad, CA, USA) was added, and the mixture was incubated in the dark for 60 min. The samples were then washed three times with PBS, decanted, resuspended in 500 μL PBS, and stained with 5 μL of 1 mg mL^−1^ calcofluor white (ICN Biomedicals Inc., Aurora, OH, USA) immediately before microscopy. Images were captured under an IX81 fluorescence microscope (Evident, Tokyo, Japan).

### 2.6 Quantitative real-time polymerase chain reaction

Total RNA was extracted from mycelia at the end of the lab-scale fermentation using a NucleoSpin RNA Plus kit (Macherey-Nagel GmbH & Co. KG, Düren, Germany) following the manufacturer’s instructions. cDNA was prepared using ReverTra Ace qPCR RT Master Mix with gDNA Remover (Toyobo, Osaka, Japan). Quantitative real-time polymerase chain reaction (PCR) was performed using Thunderbird Next SYBR qPCR Mix (Toyobo) on a MiniOpticon Detector (Bio-Rad Laboratories, Hercules, CA, USA) with the primers listed in Table S2. The transcript levels were normalized to that of histone H2B.

### 2.7 Computational fluid dynamics

CFD analyses of the 200-L and 3000-L fermentations were implemented in Ansys Fluent as described by Susukida et al. (2025). The simulation conditions are summarized in Table S3. The liquid phase was modeled as a non-Newtonian power-law fluid, with rheological parameters determined by viscometry as described by Susukida et al. (2025). Physicochemical *k_L_a* was calculated following the theoretical correlation proposed by Kawase et al. (1987).

As the scale-up criterion, *P_gV_* was selected and maintained constant across pilot scales. To estimate appropriate agitation conditions prior to pilot-scale experiments, CFD simulations were used to establish the relationship between agitation speed and *P_gV_*. For each scale (200 L and 3000 L), at least three agitation conditions were simulated, the resulting *P_gV_* values were evaluated (Table S4), and an operating condition corresponding to approximately 3 kW m^−3^ was selected. This value represents a moderate power input suitable for gas–liquid dispersion systems (Edwards et al., 1992).

To validate the accuracy of the CFD results, the uncertainty error (*E*) between the measured (see subsection 2.3) and simulated *P_gV_* was calculated using Equation 1 (Coleman and Stern, 1997):

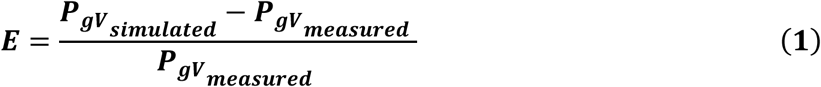

## 3 Results

### 3.1 Reduced wall growth and increased enzyme production in RolA-deficient strains

The wall growth in Sakaguchi flasks at 72 h is shown in Figure 1A. Extensive wall growth was observed in the AGΔ-GAGΔ strain, particularly where splashed culture medium adhered to the flask surface and was not readily removed by shaking-induced flow, but wall growth was markedly reduced in the AGΔ-GAGΔ-Δ*rolA* strain (Figure 1A). Quantification of biomass attached to both the spherical body and neck of the flask revealed that wall-growth dry weight was 32% lower in the AGΔ-GAGΔ-Δ*rolA* strain than in the AGΔ-GAGΔ strain (Figure 1B).

**Figure 1.**
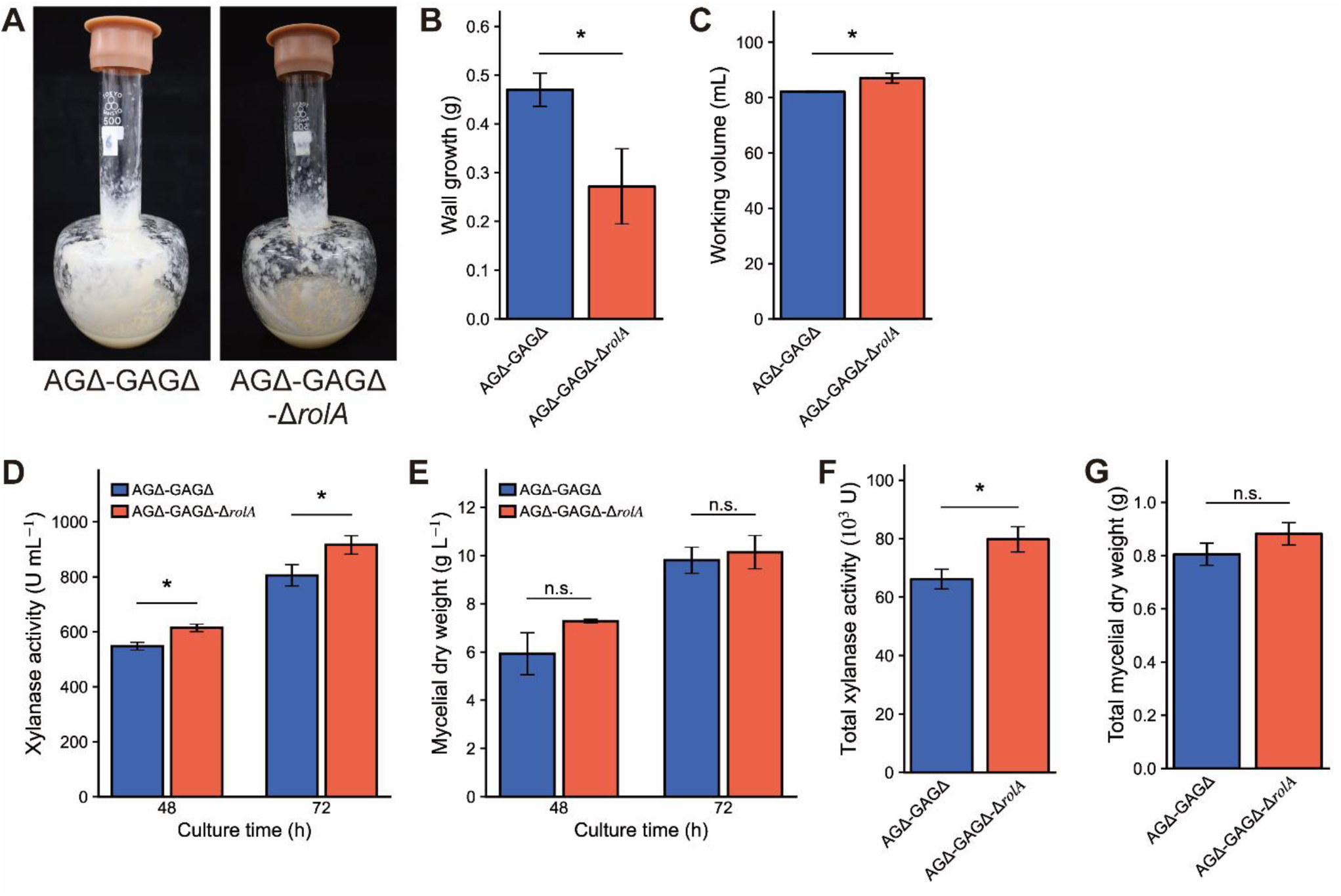
Flask-scale fermentation of AGΔ-GAGΔ and AGΔ-GAGΔ-Δ*rolA* strains. **(A)** Wall growth at the end of Sakaguchi flask fermentation. **(B)** Dry weight of wall growth at 72 h. **(C)** Working volume of culture liquid at 72 h. **(D)** Xylanase activity in culture supernatant. **(E)** Mycelial dry weight in the culture broth. **(F)** Total xylanase activity at 72 h, which is the product of the working volume and xylanase activity. **(G)** Total mycelial dry weight at 72 h. Error bars represent the standard error of the mean of three replicates. Significant differences (Student’s *t*-test): \**p* < 0.05; n.s., not significant.

The recovered liquid volume (whole culture broth) of the AGΔ-GAGΔ-Δ*rolA* was 1.1× that of the AGΔ-GAGΔ strain (Figure 1C). Xylanase activity in culture broth was up to 13% greater in the AGΔ-GAGΔ-Δ*rolA* strain than in the AGΔ-GAGΔ strain (Figure 1D); no significant difference in biomass concentration was observed between the strains (Figure 1E). Total enzyme activity was 16% greater and total biomass was 12% greater in the AGΔ-GAGΔ-Δ*rolA* strain than in the AGΔ-GAGΔ strain (Figures 1F, G).

The data for flask fermentation of the WT and WT-Δ*rolA* strains are shown in Figure S1. At 48 h, mycelial dry weight and xylanase activity were significantly higher (1.4× and 1.26×, respectively) in the WT-Δ*rolA* strain than in the WT strain. At 72 h, no statistically significant differences were observed at α = 0.05, but xylanase activity of the WT-Δ*rolA* strain remained 1.1× that of the WT strain (Figure S1).

The effects of the *rolA* gene disruption were further evaluated in a lab-scale bioreactor. The wall growth at the end of fermentation is shown in Figure 2A. Differences in biomass attachment were observed on baffles, around the impeller shaft, and on the bioreactor headspace surfaces. The dry weight of the wall growth was 20% lower in the AGΔ-GAGΔ-Δ*rolA* than in the AGΔ-GAGΔ strain (Figure 2B). Although variability between replicates was slightly higher in bioreactor cultures than in flask cultures, a similar trend was observed for culture liquid recovery: the effective working volume was higher by approximately 5% in the AGΔ-GAGΔ-Δ*rolA* strain (Figure 2C). The AGΔ-GAGΔ-Δ*rolA* strain had faster initial growth, with mycelial dry weight reaching 1.32× that of the AGΔ-GAGΔ strain at 24 h, but no significant difference was observed between the strains from 48 to 120 h (Figure 2D).

**Figure 2.**
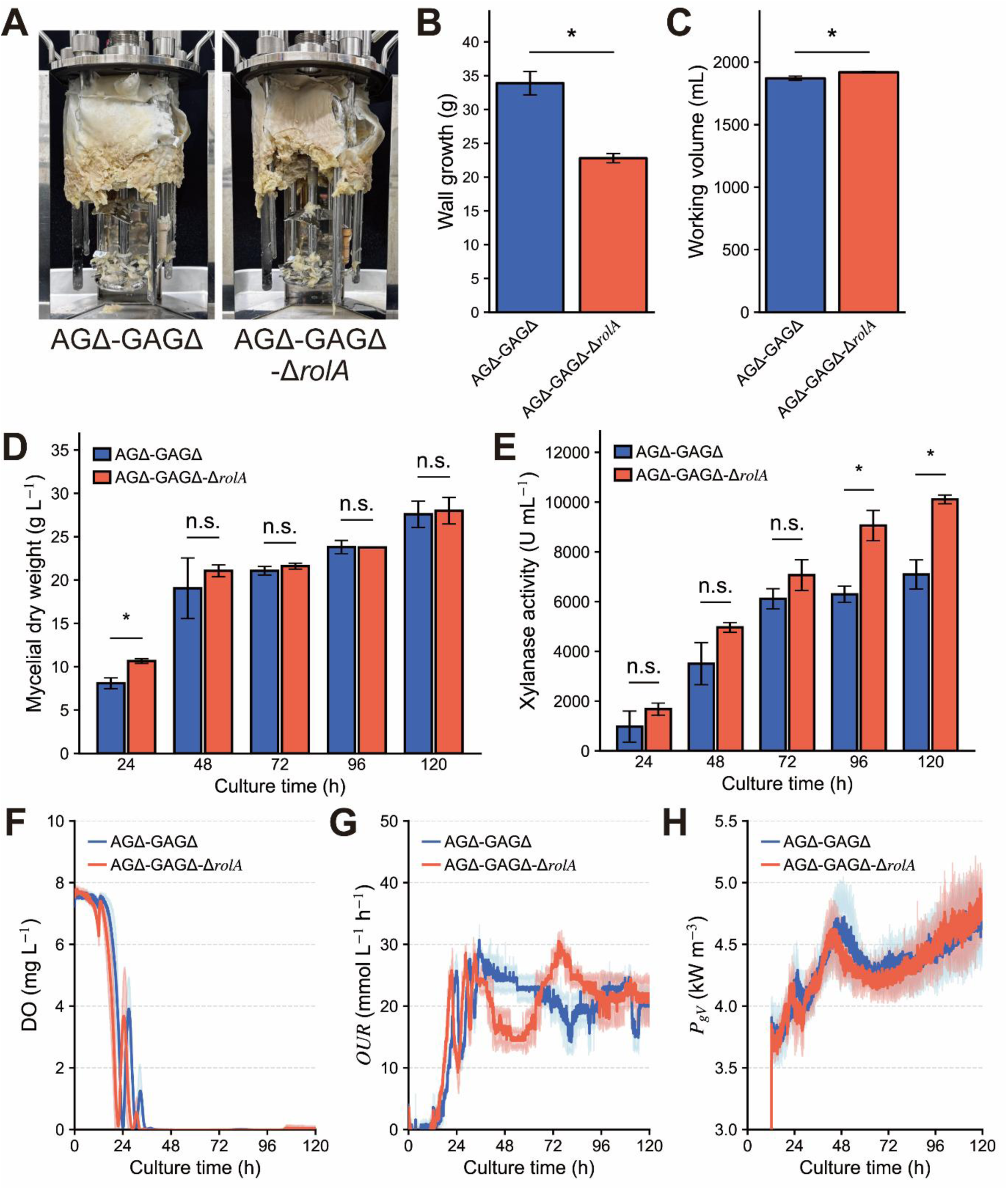
Lab-scale fermentation of AGΔ-GAGΔ and AGΔ-GAGΔ-Δ*rolA* strains. **(A)** Wall growth at 120 h of fermentation in a 4-L bioreactor. **(B)** Dry weight of wall growth at 120 h. **(C)** Working volume of culture liquid at 120 h. **(D)** Mycelial dry weight of culture liquid. **(E)** Xylanase activity in culture supernatant. **(F)** Concentration of dissolved oxygen (DO). **(G)** Oxygen uptake rate (*OUR*). **(H)** Volumetric power consumption under aeration (*P_gV_*). Error bars in **(B**–**E)** represent the standard error of the mean among three replicates. Shades in **(F**–**H)** represent the standard deviation of the mean among three replicates. Significant differences (Student’s *t*-test): \**p* < 0.05; n.s., not significant.

Xylanase activity in the bioreactor culture broth was consistently higher in the AGΔ-GAGΔ-Δ*rolA* strain throughout fermentation and reached 1.44× and 1.43× that of the AGΔ-GAGΔ strain at 96 h and 120 h, respectively (Figure 2E). In the AGΔ-GAGΔ-Δ*rolA* strain, DO was depleted by 21 h, followed by a transient spike and then zero until the end of fermentation (Figure 2F). A similar trend was observed in the AGΔ-GAGΔ-Δ*rolA* strain, although DO depletion occurred approximately 3 h later. The *OUR* increased earlier in the AGΔ-GAGΔ-Δ*rolA* strain (Figure 2G). It was lower in the AGΔ-GAGΔ-Δ*rolA* strain than in the AGΔ-GAGΔ strain between 36 and 66 h, but was higher between 66 and 96 h (Figure 2G). Under aerated conditions, agitation power consumption was similar between the strains, although the AGΔ-GAGΔ-Δ*rolA* strain tended to require slightly lower power input up to 96 h (Figure 2H).

### 3.2 RolA expression on the cell surface of the AGΔ-GAGΔ-Δ*rolA* strain during stirred-tank bioreactor fermentation

Quantitative analysis of *rolA* expression confirmed that the *rolA* transcript in the AGΔ-GAGΔ-Δ*rolA* strain was nearly abolished by gene disruption (Figure 3A). The expression level in the AGΔ-GAGΔ strain increased until the end of fermentation; at 120 h, it was 9.8× that at 24 h. Fluorescence microscopy revealed that RolA was expressed on the surface of the AGΔ-GAGΔ hyphae in the liquid fermentation condition, whereas no fluorescence signal was observed in the AGΔ-GAGΔ-Δ*rolA* strain (Figure 3B).

**Figure 3.**
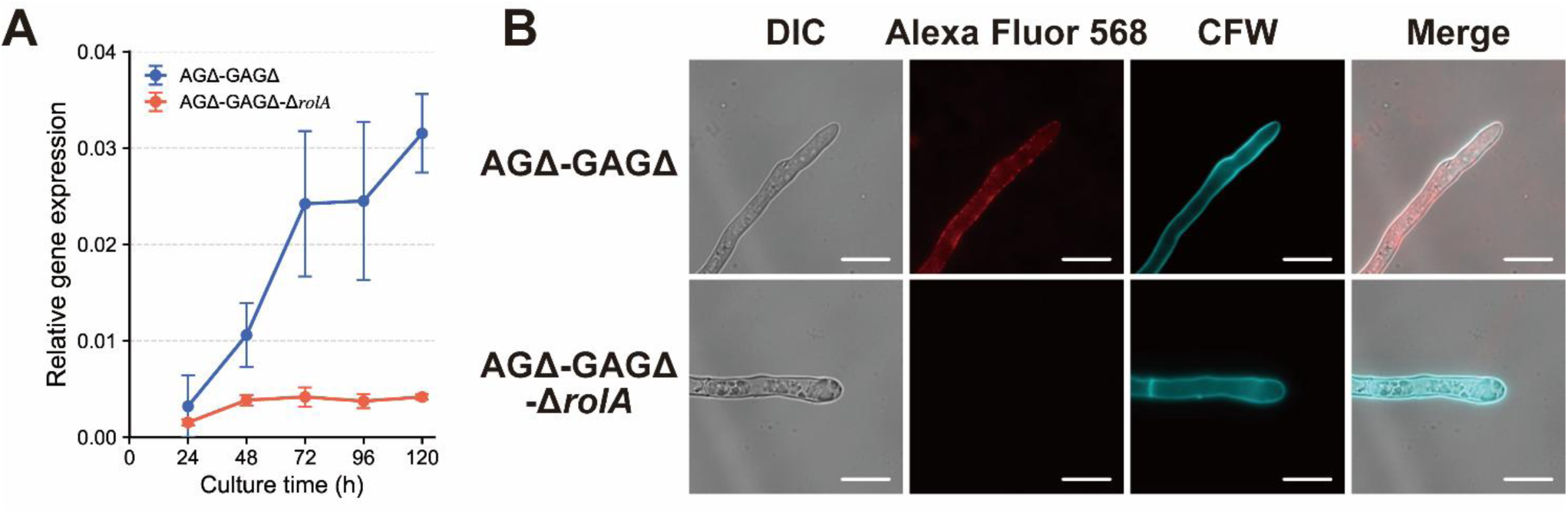
Expression of *rolA* gene and cell-surface RolA in AGΔ-GAGΔ and AGΔ-GAGΔ-Δ*rolA* strains in lab-scale bioreactor fermentation. **(A)** Quantitative PCR analysis of the *rolA* gene of hyphae. *Y*-axis values were normalized to histone h2b expression, which was considered as 1. **(B)** Immuno-fluorescent labeling of cell-surface RolA on hyphae at 120 h. DIC, differential interference contrast microscopy; CFW, calcofluor white. Scale bars: 10 μm.

### 3.3 Improved hydrodynamic properties of the AGΔ-GAGΔ-Δ*rolA* strain in pilot-scale fermentation

Viscometry showed similar power-law behavior between the AGΔ-GAGΔ and AGΔ-GAGΔ-Δ*rolA* strains (Figure S2). Scale-up evaluation of the AGΔ-GAGΔ-Δ*rolA* strain was performed using constant *P_gV_* (3 kW m^−3^). Preliminary CFD analyses using experimentally measured broth viscosity estimated agitation speeds corresponding to approximately 3 kW m^−3^ to be 400 rpm for the AGΔ-GAGΔ strain and 390 rpm for the AGΔ-GAGΔ-Δ*rolA* strain at the 200-L scale, and 263 rpm for the AGΔ-GAGΔ-Δ*rolA* strain at the 3000-L scale (Table S3).

Pilot-scale fermentation was conducted only for the AGΔ-GAGΔ-Δ*rolA* strain under the agitation conditions determined by CFD analysis. Mycelial dry weight and xylanase activity were nearly identical between the 200-L and 3000-L scales up to 96 h of fermentation (Figures 4A, B). At later time points, mycelial dry weight was slightly higher at the 3000-L scale than at the 200-L scale (Figure 4A), whereas xylanase activity was higher at the 200-L scale, reaching 1.15× that of the 3000-L scale at 168 h (Figure 4B). The overall DO profiles were similar between the two scales, although DO was depleted approximately 3 h earlier in the 200-L culture (Figure 4).

**Figure 4.**
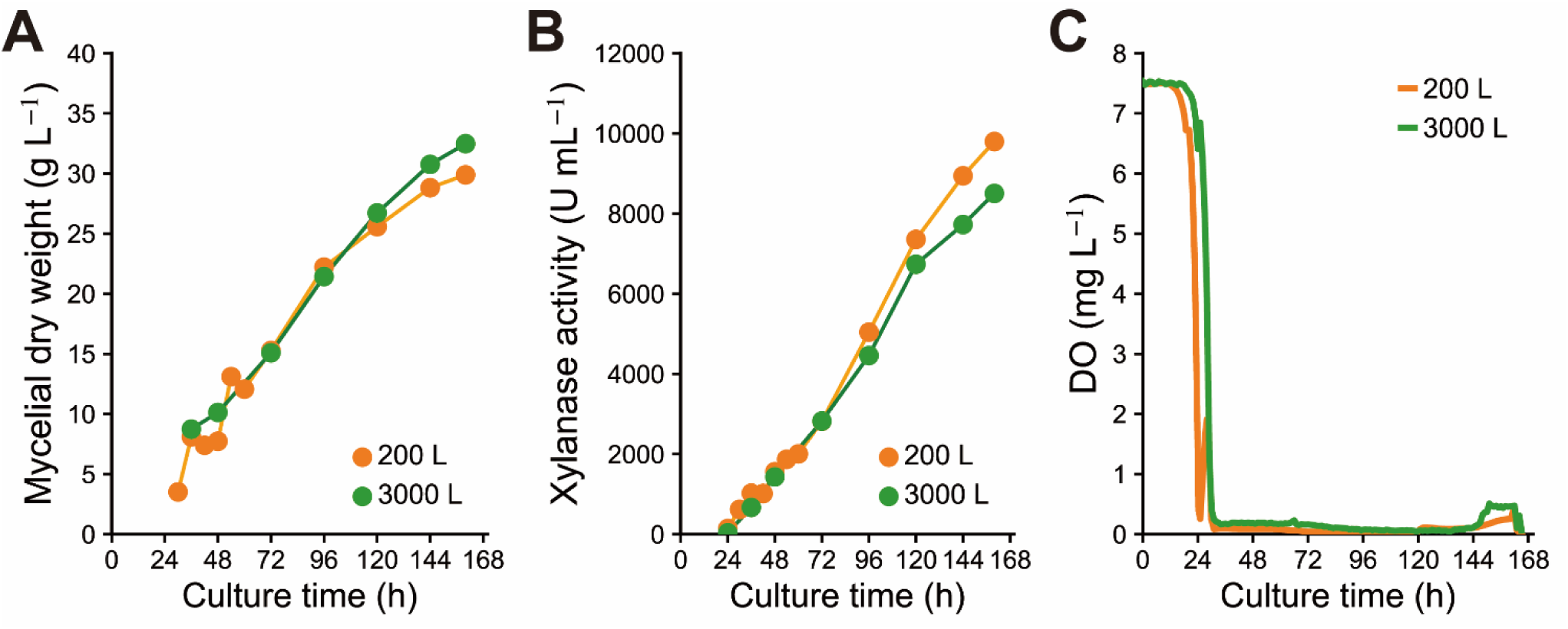
Pilot-scale fermentation of AGΔ-GAGΔ-Δ*rolA* strain. **(A)** Mycelial dry weight. **(B)** Xylanase activity in culture supernatant. **(C)** Dissolved oxygen (DO).

CFD analysis results are shown in Figure 5. At the 200-L scale, flow patterns were markedly improved in the AGΔ-GAGΔ-Δ*rolA* strain in comparison with the AGΔ-GAGΔ strain as pathline expands towards the edge of the bioreactor in AGΔ-GAGΔ-Δ*rolA* strain (Figure 5A). Fluid circulation and inter-impeller flow exchange were also enhanced in AGΔ-GAGΔ-Δ*rolA* (Figure 5B), resulting in a 1.35× increase in the average velocity and a 1.22× increase in the average shear rate in comparison with those of AGΔ-GAGΔ. Considerable gas accumulation was observed near the liquid surface around the impeller shaft in the AGΔ-GAGΔ cultures but not in the AGΔ-GAGΔ-Δ*rolA* cultures (Figure 5C). The reactor-averaged *k_L_a* of AGΔ-GAGΔ-Δ*rolA* was 92% that of AGΔ-GAGΔ and the average shear stress was 76% of that of AGΔ-GAGΔ.

**Figure 5.**
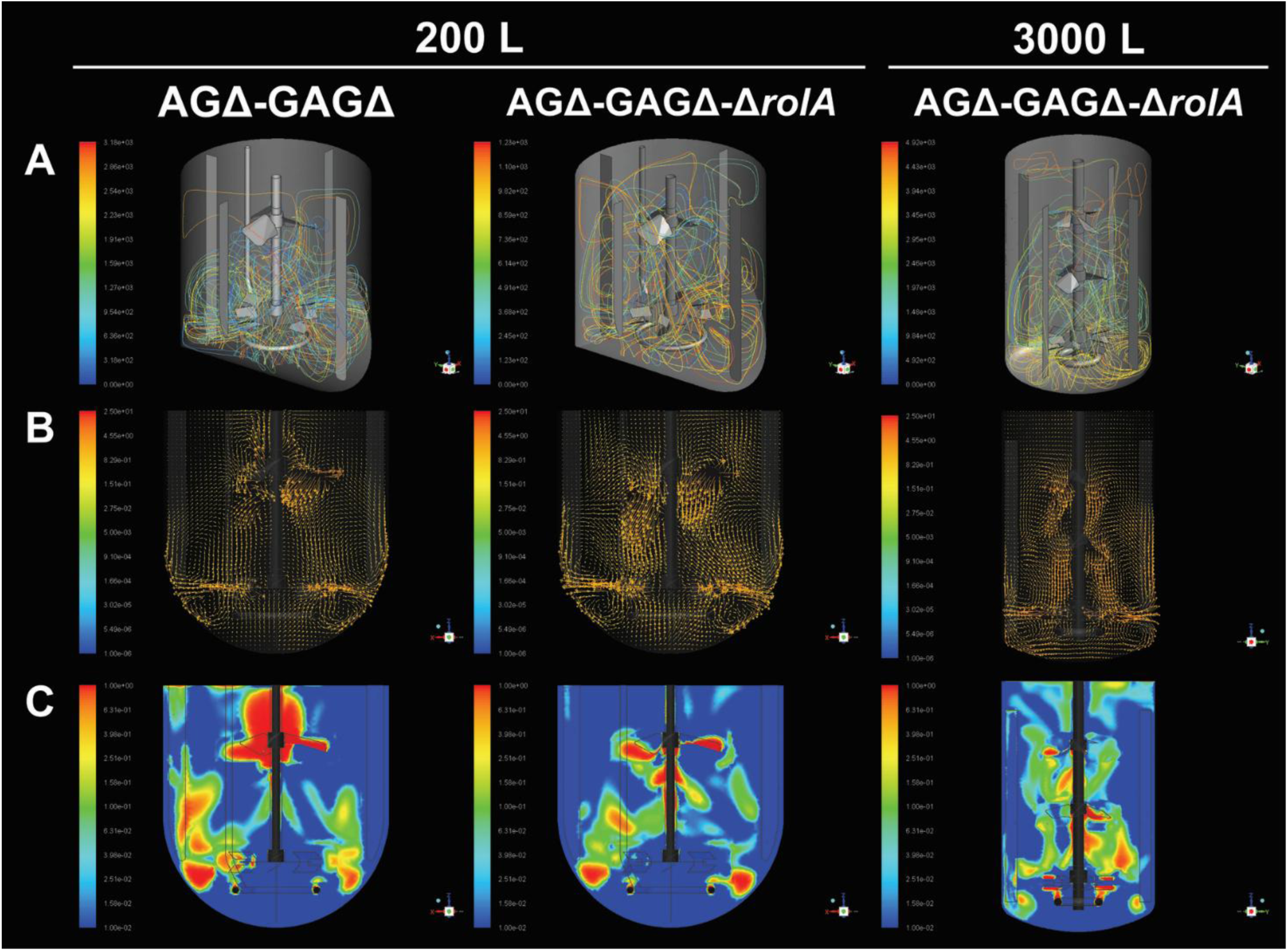
Computational fluid dynamics analysis of pilot-scale fermentation of AGΔ-GAGΔ and AGΔ-GAGΔ-Δ*rolA* strains. AGΔ-GAGΔ culture (400 rpm) and AGΔ-GAGΔ-Δ*rolA* culture (390 rpm) at the 200-L scale and AGΔ-GAGΔ-Δ*rolA* culture (263 rpm) at the 3000-L scale. **(A)** Pathline visualization of three-dimensional flow. Different colors denote different streamlines from impellers. **(B)** Velocity vectors. **(C)** Gas fraction contours. Color scales in **(B)** and **(C)** denote the magnitude of each parameter (red, highest; blue, lowest values).

CFD analysis at the 3000-L scale was conducted only for the AGΔ-GAGΔ-Δ*rolA* strain. The flow pattern showed well-developed circulation extending to the upper region of the bioreactor (Figure 5A). Velocity vector distributions were also well developed compared with the 200-L AGΔ-GAGΔ-Δ*rolA* simulation, and no considerable gas accumulation around the shaft was observed. The *k_L_a* values for the AGΔ-GAGΔ-Δ*rolA* strain were nearly identical at both scales, whereas average shear rate and liquid velocity were 0.61× and 1.68×, respectively, those at the 200-L scale.

Additional CFD simulations at different agitation speeds demonstrated consistent relationships among impeller tip speed (*V_tip_*), *P_gV_*, *k_L_a*, and shear stress across scales (Figure S6, Table S4). When all CFD conditions were considered together, the volume-averaged *k_L_a* showed a strong power-law relationship with *P_gV_* (Figure S6C):

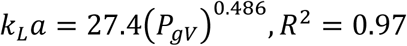

While *k_L_a* showed similar dependence on *P_gV_* regardless of scale, the 3000-L condition exhibited higher average shear stress at comparable *P_gV_* values to that under the 200-L condition.

The uncertainty of CFD results was examined by comparing simulated and measured *P_gV_* values. At the 200-L scale, the measured *P_gV_* value at the end of fermentation of the AGΔ-GAGΔ strain was 3.57 kW m^−3^ and the simulated one was 3.33 kW m^−3^, corresponding to an uncertainty error *E* (Eq. 1) of 6.7%. At the 3000-L scale, the measured *P_gV_* value for the AGΔ-GAGΔ-Δ*rolA* strain at 168 h was 3.30 kW m^−3^ and the simulated value was 3.21 kW m^−3^, giving an *E* of 2.7%.

## 4 Discussion

Although hydrophobins are classically associated with aerial hyphae, conidia, and fruiting bodies (Wessels, 1997; Wosten, 2001), several studies have demonstrated that hydrophobins are also expressed during liquid fermentation and can be detected in the fungal cell wall, at the hyphal surface, and in the culture medium (Nakari-Setälä et al., 1996; de Vries et al., 1999; Lacroix et al., 2008). In the present study, *rolA* transcript levels remained low throughout lab-scale fermentation of the AGΔ-GAGΔ strain (Figure 3A). Nevertheless, the *rolA* expression level increased during lab-scale bioreactor fermentation, and immunofluorescence microscopy clearly demonstrated the presence of RolA on the hyphal surface of the AGΔ-GAGΔ strain (Figure 3B). These observations indicate that even low levels of *rolA* expression are sufficient to maintain surface-associated hydrophobin during liquid fermentation. Therefore, RolA may still influence the physicochemical properties of fungal hyphae despite the low transcriptional activity of its gene.

The absence of RolA reduced wall growth and increased working volume across fermentation scales (Figures 1 and 2). This result is consistent with the established role of hydrophobins in increasing fungal surface hydrophobicity and promoting adhesion at hydrophilic–hydrophobic interfaces (Wessels, 1997; Takahashi et al., 2005). Class I hydrophobins, such as SC3 in *Schizophyllum commune*, RodA in *Aspergillus fumigatus*, and RolA in *A. oryzae*, self-assemble into amphipathic films and tightly packed amyloid fibrils known as rodlets that enable fungal attachment to a variety of surfaces and confer surface-associated functions (Wang et al., 2004; Linder et al., 2005; Morris et al., 2011; Terauchi et al., 2020). Consistent with these data, disruption of *rolA* in the AGΔ-GAGΔ strain weakened the interaction between fungal biomass and bioreactor surfaces, reducing wall growth and improving culture recovery, as also reported in *Trichoderma reesei* (Bailey et al., 2002; Paulsen et al., 2014).

Conidial hydrophobicity is lower in hydrophobin-deficient strains than in their parental strains because the hydrophobic rodlet layer is absent from the cell surface (Whiteford and Spanu, 2001; Paris et al., 2003; Askolin, 2006; Ida et al., 2026). Deletion of specific hydrophobins in *Trichoderma* species eliminates the lag phase before spore germination and accelerates germination (Cai et al., 2021). In this study, both WT-Δ*rolA* and AGΔ-GAGΔ-Δ*rolA* strains had faster initial growth compared to parental strains, as indicated by earlier biomass accumulation in flask-scale fermentation (Figures 1 and S1). The AGΔ-GAGΔ-Δ*rolA* strain also showed faster initial biomass accumulation and DO depletion than its parental strain in lab-scale fermentation (Figure 2). These results suggest that disruption of *rolA* promoted earlier establishment of growth in liquid culture, thereby shortening the time required to reach productive fermentation. From a bioprocessing perspective, such accelerated culture development is advantageous because it can increase the overall process productivity while reducing fermentation time.

Previous studies of hydrophobin-deficient fungi have primarily focused on improvements in foam control during fermentation and downstream processing, which facilitate industrial fermentation. Hydrophobins are highly surface-active proteins that accumulate at gas–liquid interfaces, stabilize bubbles, and in some cases form highly elastic or rigid interfacial membranes (Wosten et al., 1994). Although the ability to stabilize foams and emulsions with hydrophobins makes them attractive biosurfactants for industrial application, several studies and patents have proposed hydrophobin-deficient strains as advantageous for liquid fermentation due to their reduced tendency to generate persistent foam (Nakari-Setälä et al., 2001; Bailey et al., 2002; Hektor and Scholtmeijer, 2005; Paulsen et al., 2014). However, the consequences of hydrophobin deficiency for bulk broth rheology and bioreactor hydrodynamics have not been investigated.

Foaming control has long been recognized as an important aspect of industrial fermentation because excessive foam can reduce operational efficiency. Adsorption of proteins and peptides at gas– liquid interfaces suppresses bubble coalescence and promotes the formation of persistent foams (Prins and van’t Riet, 1987). Fungal biosurfactants that stabilize foams include hydrophobins, repellents, fungispumins, and cerato-platanins (Razafindralambo et al., 1996; Kershaw and Talbot, 1998; Zapf et al., 2007; Frischmann et al., 2013). Antifoaming agents are widely used to destabilize interfacial films and prevent excessive foam formation. However, the same interfacial phenomena that govern foam stability also influence gas–liquid mass transfer. Stabilization of liquid films surrounding bubbles suppresses bubble coalescence, resulting in smaller bubbles and a larger gas–liquid interfacial area, whereas antifoaming agents promote bubble coalescence and may reduce the interfacial area available for oxygen mass transfer (Prins and van’t Riet, 1987; Vardar-Sukan, 1998). Thus, hydrophobins may influence not only foam formation but also the hydrodynamic and oxygen-transfer characteristics of liquid cultures.

Interestingly, in comparison with the AGΔ-GAGΔ strain, the AGΔ-GAGΔ-Δ*rolA* strain had altered hydrodynamic behavior despite only minor differences in *OUR* of experimental measurements and *k_L_a* of CFD analysis (Figure 2, Table 1); marked differences between the two strains were observed in flow patterns, gas distribution, and circulation behavior in 200-L CFD (Figures 5, S3, and S4). One possible explanation is that the absence of RolA reduced bubble stabilization in the bioreactor. Because RolA adsorbs strongly to gas–liquid interfaces and can stabilize bubbles and foams, the AGΔ-GAGΔ strain may have generated more stable gas accumulation despite similar overall oxygen transfer rates in both strains. Consistent with this hypothesis, CFD simulations revealed greater gas accumulation around the impeller shaft in the AGΔ-GAGΔ culture than in the AGΔ-GAGΔ-Δ*rolA* culture (Figure 5). Thus, hydrophobin deficiency may improve hydrodynamics without necessarily increasing vessel-averaged *k_L_a*. CFD analysis performed at the same *P_gV_* demonstrated stronger liquid circulation, improved inter-impeller exchange, and more uniform flow patterns in the AGΔ-GAGΔ-Δ*rolA* culture (Figure 5). Although the average *k_L_a* was slightly reduced, average shear stress also decreased and local shear stress distribution was milder in AGΔ-GAGΔ-Δ*rolA* (Figures S4, Table 1), suggesting reduced spatial heterogeneity within the vessel. Gas accumulation around impellers under aeration typically decreases their pumping capacity, especially in highly viscous liquids such as fungal broth, hence suppressing the overall circulation within the bioreactor (Susukida et al., 2025). These results indicate that the improvement in fermentation performance in the absence of RolA was not driven solely by oxygen transfer, but rather by a combination of reduced wall growth, altered rheology, improved circulation, and more homogeneous hydrodynamic conditions.

**Table 1.**
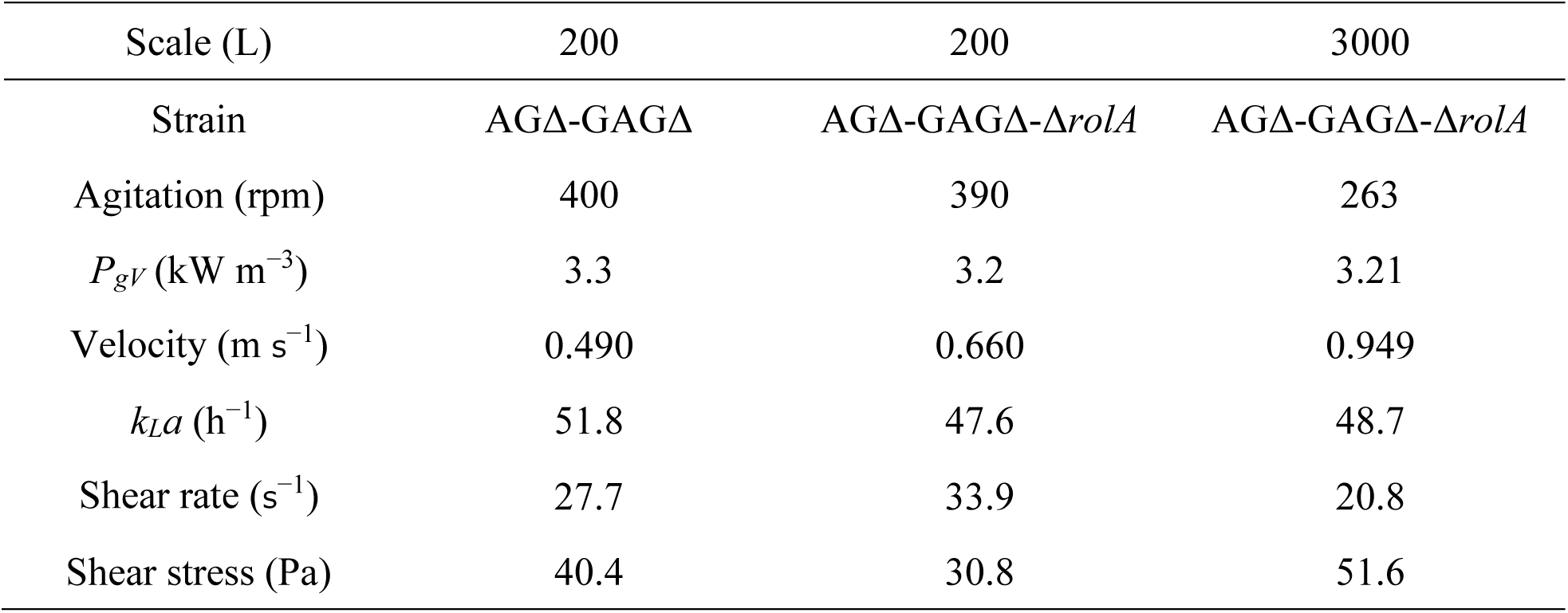
Results of computational fluid dynamics analysis.

Aerobic fermentation is commonly scaled up using criteria such as power consumption, oxygen mass transfer, and agitation parameters (Garcia-Ochoa and Gomez, 2009). Because oxygen mass transfer and mixing are among the major limitations in filamentous fungal fermentation, we selected *P_gV_* as the scale-up criterion. CFD simulations demonstrated a strong correlation between *P_gV_* and *k_L_a* across several conditions (Figure S6), supporting the use of *P_gV_* as a practical scale-up parameter in this system.

Although comparable *P_gV_* values showed similar *k_L_a* values across scales, CFD analysis revealed that average shear stress increased with up-scaling (Figure S6D); this behavior resulted from the higher *V_tip_* required in a larger bioreactor to maintain the target power input. Consequently, the culture broth was exposed to higher local shear rates (Tables 1, S4, Figure S6D), potentially amplifying differences in shear-thinning behavior between strains. Under these conditions, the rheological advantages of the AGΔ-GAGΔ-Δ*rolA* strain become more apparent, contributing to the improved circulation characteristics suggested by the CFD analysis (Figure 5). Despite higher average shear at the 3000-L scale, the AGΔ-GAGΔ-Δ*rolA* strain maintained dry mycelial weight and enzyme activity comparable to those at the 200-L scale (Figure 4). These results suggest that the benefits of *rolA* deletion were retained under industrially relevant mixing conditions and may become increasingly advantageous as the fermentation scale increases.

The CFD predictions were validated by experimentally measuring power consumption, yielding an uncertainty error of 6.7% at the 200-L scale and 2.7% at the 3000-L scale, both of which were within the range reported in a previous CFD validation study (Sadino-Riquelme et al., 2022). This validation supports the reliability of the hydrodynamic comparisons used throughout this study. Together, the CFD and fermentation results demonstrated that the *P_gV_* scale-up strategy applied to AGΔ-GAGΔ-Δ*rolA* cultures successfully maintained fermentation performance and provide quantitative insight into the hydrodynamic consequences of strain-dependent rheological differences.

The contribution of hydrophobins to bulk culture rheology has rarely been discussed despite decades of research on these proteins One possible explanation is that wild-type *Aspergillus* strains possess dominant aggregation factors, α-1,3-glucan and galactosaminogalactan, whose effects on viscosity and macro-hydrodynamics may mask the smaller contribution of hydrophobins (Miyazawa et al., 2019; Susukida et al., 2025). Comparative analysis of the rheological properties of the AGΔ-GAGΔ and AGΔ-GAGΔ-Δ*rolA* strains in the present study reveals this previously overlooked contribution.

Further studies are required to determine the mechanisms linking hydrophobins to rheology and hydrodynamics. Possible factors include their effects on rheological properties of both micro- and macro-scale mycelial suspensions, bubble size distribution, and gas–liquid interfacial behavior. To our knowledge, this study provides the first evidence that hydrophobin deficiency can influence not only wall growth, but also culture rheology and bioreactor hydrodynamics during large-scale filamentous fungal liquid fermentation.

## 5 Conclusion

In this study, we addressed major challenges in filamentous fungal liquid fermentation, namely control of mycelial morphology, increased broth viscosity, and wall growth, through strain engineering. The previously developed AGΔ-GAGΔ strain had dispersed growth, reduced broth viscosity, and improved enzyme productivity compared with the wild-type strain. In the present study, we further constructed the AGΔ-GAGΔ-Δ*rolA* strain by disrupting the *rolA* gene encoding a hydrophobin. This strain showed reduced wall growth in both flask and lab-scale fermentations, increased effective working volume, and improved recovery of recombinant enzyme activity and biomass. Pilot-scale fermentations demonstrated successful scale-up to 200-L and 3000-L bioreactors while maintaining enzyme productivity and improved hydrodynamics.

Combining hydrophobin deficiency with a hyphal-dispersion background is an effective strategy for improving filamentous fungal liquid fermentation, and our results suggest that AGΔ-GAGΔ-Δ*rolA* is a promising strain for industrial enzyme production.

## Data availability statement

The original contributions presented in the study are included in the article and Supplementary Material; further inquiries can be directed to the corresponding authors.

## Author contributions

SS: Conceptualization, Data curation, Investigation, Methodology, Visualization, Writing–original draft, Writing–review & editing, Formal Analysis, Validation. YB: Writing–review & editing, Methodology, Investigation, MF: Writing–review & editing, Investigation, YN: Writing–review & editing, Investigation, KiM: Conceptualization, Methodology, Writing–review and editing. KeM: Writing–review and editing. AY: Writing–review and editing. YK: Methodology, Software, Visualization, Resources, Writing–review and editing. HH: Writing–review & editing, Methodology, Resources, Supervision, KA: Conceptualization, Project administration, Resources, Funding acquisition, Supervision, Writing–review and editing.

## Acknowledgements

We gratefully acknowledge the support of the Green Earth Institute, which provided the pilot-scale infrastructure used to conduct this study. We also thank Prof. Motoaki Sano (Kanazawa Institute of Technology) for providing host strains.

