## Supplementary Material for "Improved recombinant protein production and scale-up fermentation of *Aspergillus oryzae* hyphal-dispersion hydrophobin-deficient strain"

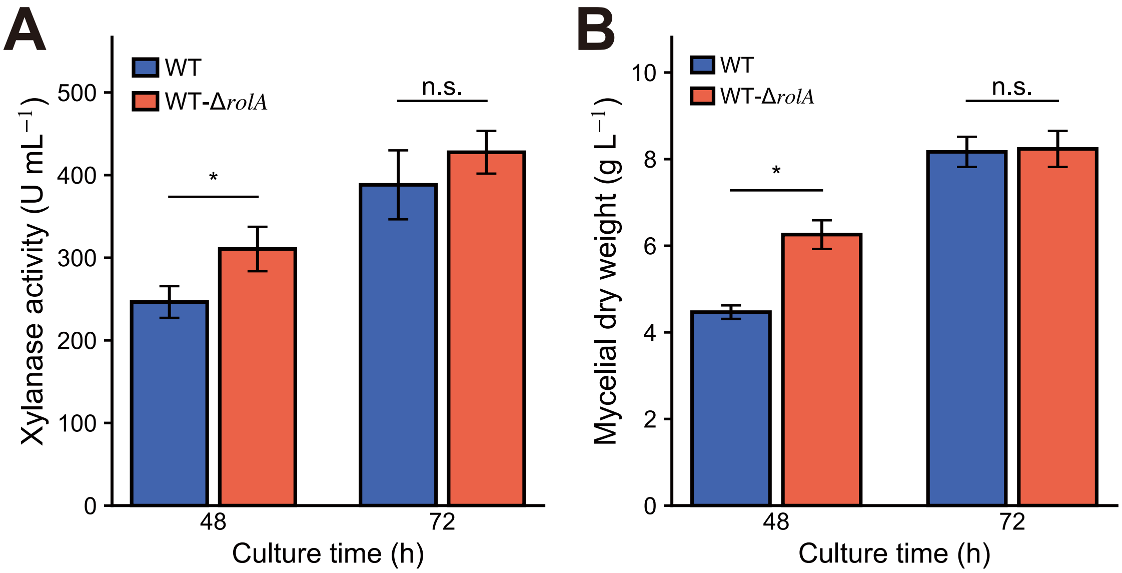


**Figure S1. Flask-scale cultivation of WT and WT-Δ*rolA* strains.**

**(A)** Dry mycelial weight in culture broth. **(B)** Xylanase activity in culture supernatant. Error bars represent standard error of the mean of three replicates. Significant differences (Student's *t*-test): $*p < 0.05$; n.s., not significant.

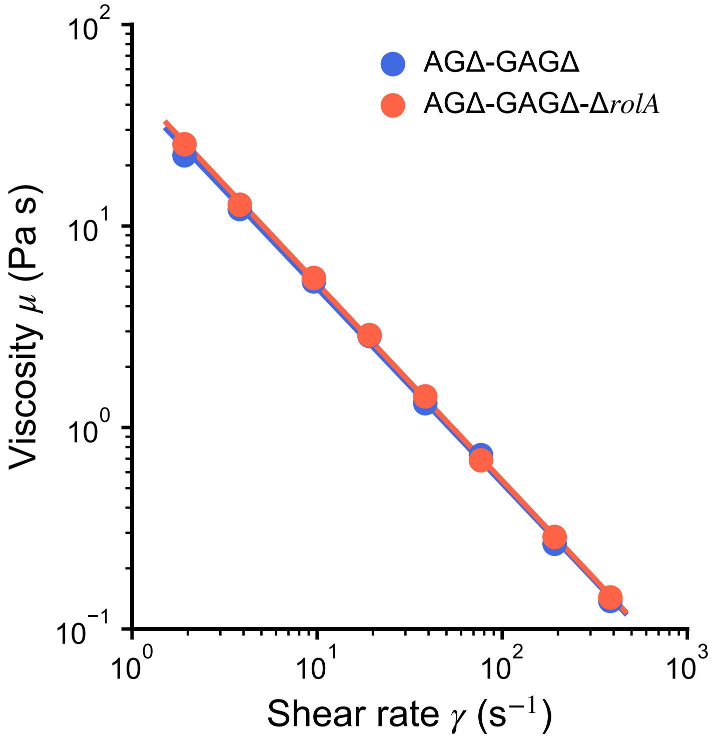


**Figure S2. Rheogram of culture broth of AGΔ-GAGΔ and AGΔ-GAGΔ-Δ*rolA* strains.**

Viscosity was measured with an E-type viscometer. Solid lines are the power trendlines of each strain.

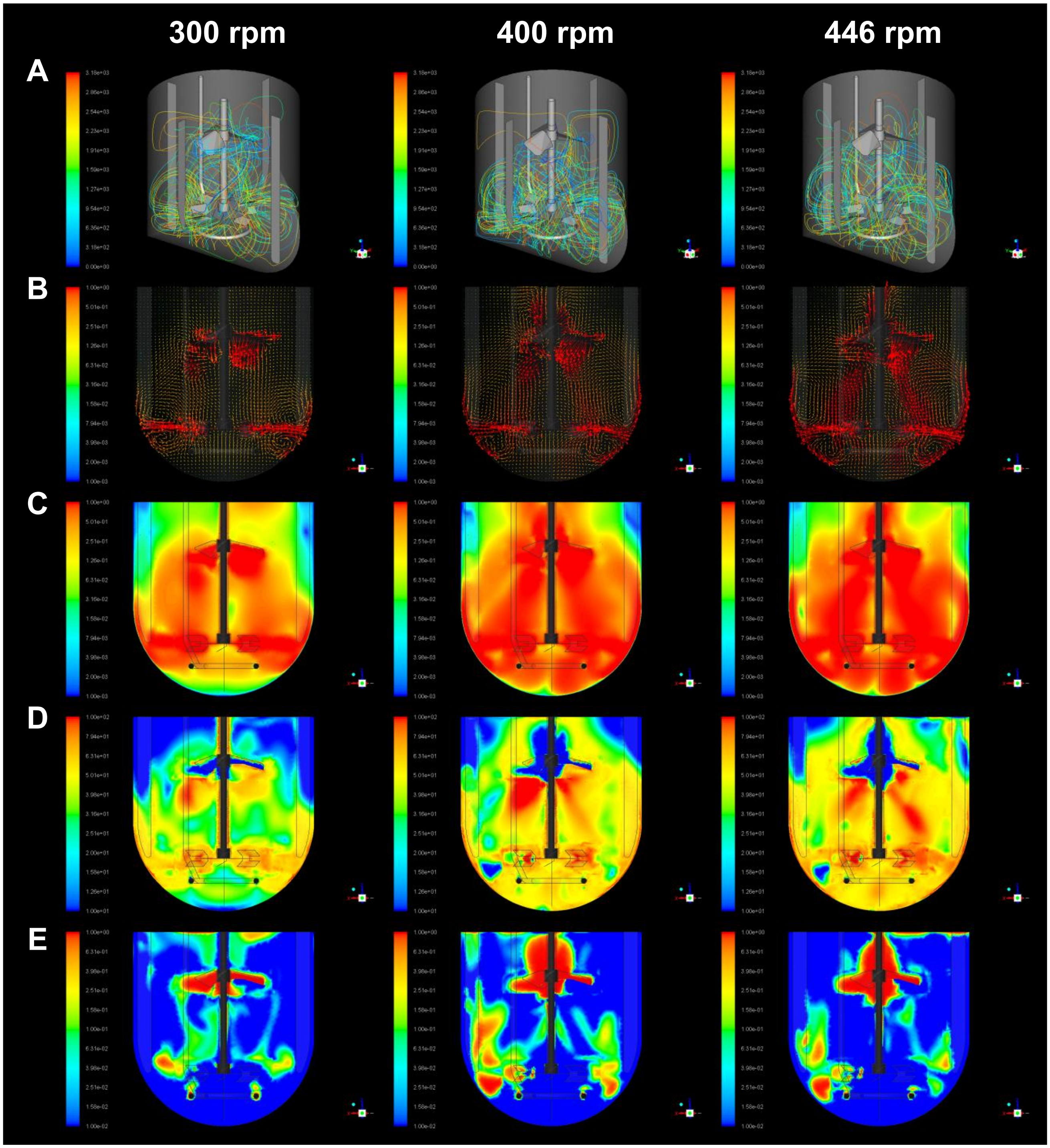


**Figure S3. Computational fluid dynamics (CFD) analysis of 200-L fermentation of AGΔ-GAGΔ strain in three agitation conditions.**

**(A)** Pathline visualization of three dimensional flow. Different colors denote different streamlines from impellers. **(B)** Velocity vectors. **(C)** Velocity contours. **(D)** Shear stress contours. **(E)** Gas fraction contours. Color scales in **(B–E)** denote the magnitude of each parameter (red, highest; blue, lowest). The results at 400 rpm are reproduced from Figure 5 for direct comparison.

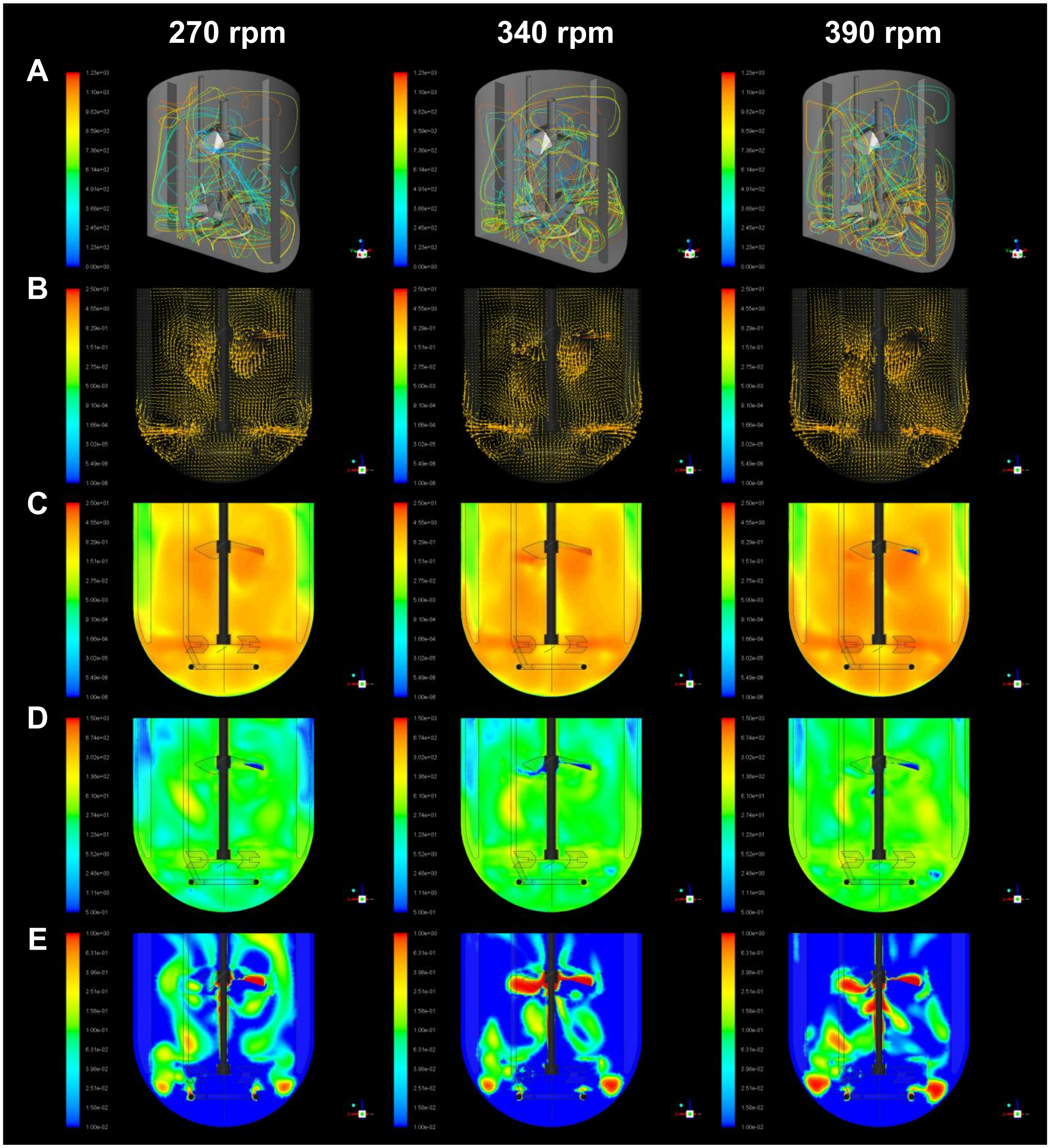


**Figure S4. CFD analysis of 200-L fermentation of AGΔ-GAGΔ-Δ*rolA* strain in three agitation conditions.**

**(A)** Pathline visualization of three-dimensional flow. Different colors denote different streamlines from impellers. **(B)** Velocity vectors. **(C)** Velocity contours. **(D)** Shear stress contours. **(E)** Gas fraction contours. Color scales in **(B–E)** denote the magnitude of each parameter (red, highest; blue, lowest). The results at 390 rpm are reproduced from Figure 5 for direct comparison.

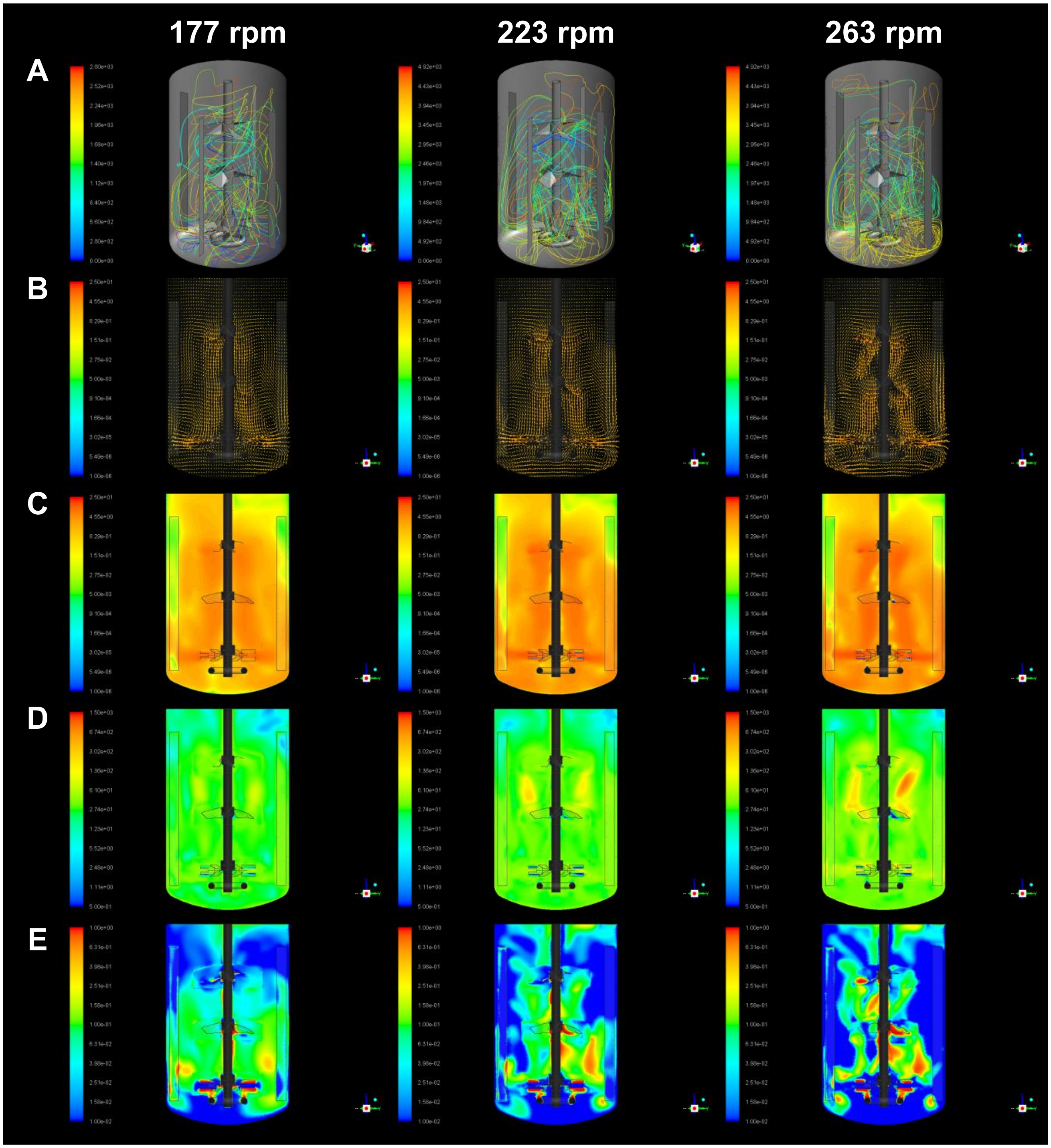


**Figure S5. CFD analysis of 3000-L fermentation of AGΔ-GAGΔ-Δ*rolA* strain in three agitation conditions.**

**(A)** Pathline visualization of three-dimensional flow. Different colors denote different streamlines from impellers. **(B)** Velocity vectors. **(C)** Velocity contours. **(D)** Shear stress contours. **(E)** Gas fraction contours. Color scales in **(B–E)** denote the magnitude of each parameter (red, highest; blue, lowest). The results at 263 rpm are reproduced from Figure 5 for direct comparison.

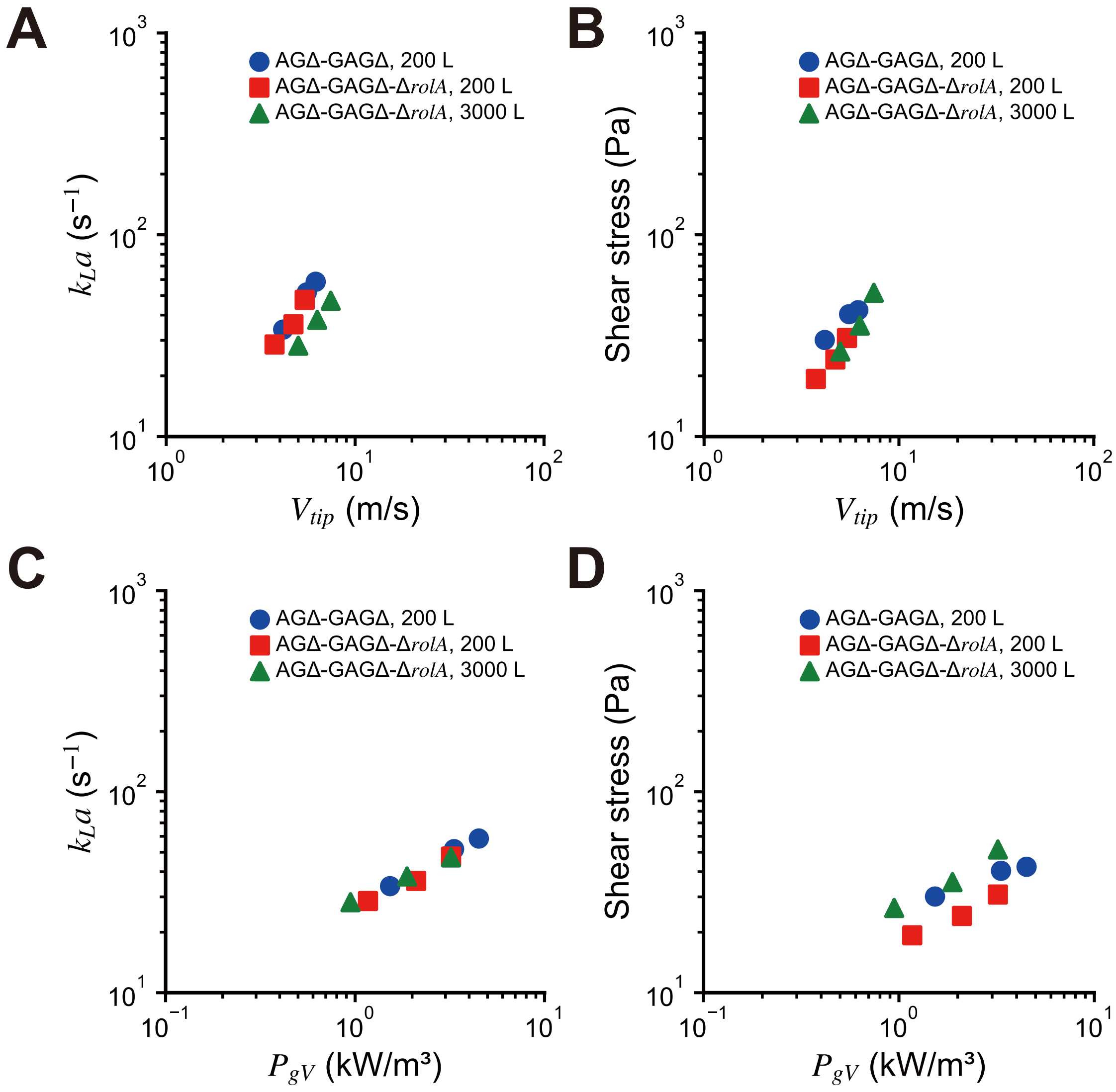


**Figure S6. Comparison of CFD results across fermentation scales.**

**(A**, **B)** Relationship between impeller tip speed ($V_{tip}$) and volume-averaged **(A)** $k_La$ or **(B)** shear stress. **(C**, **D)** Relationship between volumetric power consumption ($P_{gV}$) and volume-averaged **(C)** $k_La$ or **(D)** shear stress.

**Table S1. Strains used and their genotypes.**

| Strain | Genotype |
| --- | --- |
| WT | Δ*ligD*::*sC*, Δ*adeA*::*ptrA*, *PglaA142*-xynF1::*niaD* |
| WT-Δ*rolA* | Δ*ligD*::*sC*, Δ*adeA*::*ptrA*, Δ*rolA*::*adeA*, *PglaA142-xynF1*::*niaD* |
| AGΔ-GAGΔ | Δ*ligD*::*sC*, Δ*adeA*::*ptrA*, *agsA*::*loxP*, *agsB*::*loxP*, *agsC*::*loxP*, *sphZugeZ*::*adeA*, *PglaA142-xynF1*::*niaD* |
| AGΔ-GAGΔ-Δ*rolA* | Δ*ligD*::*sC*, Δ*adeA*::*ptrA*, *agsA*::*loxP*, *agsB*::*loxP*, *agsC*::*loxP*, *sphZugeZ*::*loxP*, Δ*rolA*::*adeA*, *PglaA142-xynF1*::*niaD* |

**Table S2. Primers used for quantitative polymerase chain reaction.**

| Oligonucleotide | Sequence (5′ to 3′) |
|---|---|
| H2b_F_qPCR | CCCTGCTGAGAAGAAGGAAG |
| H2b_R_qPCR | CGAGTGGAGATTCCAGTATC |
| RolA_F_qPCR | ATGCAGTTCTCCGTCGCCGC |
| RolA_R_qPCR | GTTGGAGAGGAGGCCAGCAA |

**Table S3. CFD analysis conditions.**

| Scale (L) | 200 | 3000 |
|---|---|---|
| Vessel diameter (mm) | 590 | 1200 |
| Liquid volume (L) | 150 | 2000 |
| Impellers | HR100/HS100 | 2×HR100/HS100 |
| Impeller diameter (mm) | 265 | 540 |
| Liquid phase density ($kg\ m^{-3}$) | 1040 | 1040 |
| Liquid phase viscosity $\mu^{\dagger}$ (Pa s) | AGΔ-GAGΔ: 45.771× $\gamma^{-0.969}$<br>AGΔ-GAGΔ-Δ*rolA*: 49.365× $\gamma^{-0.978}$ | AGΔ-GAGΔ-Δ*rolA*: 49.365× $\gamma^{-0.978}$ |
| Agitation rate (rpm) | AGΔ-GAGΔ: 300, 400, 446<br>AGΔ-GAGΔ-Δ*rolA*: 270, 340, 390 | AGΔ-GAGΔ-Δ*rolA*: 177, 223, 263 |
| Aeration ($L\ min^{-1}$) | 96 | 1440 |
| 3D model | 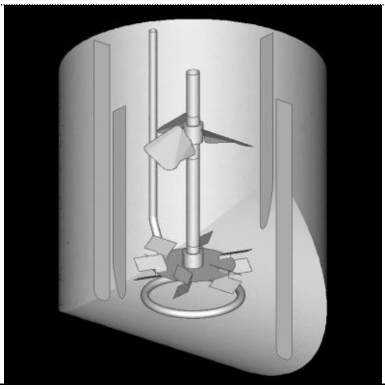 | 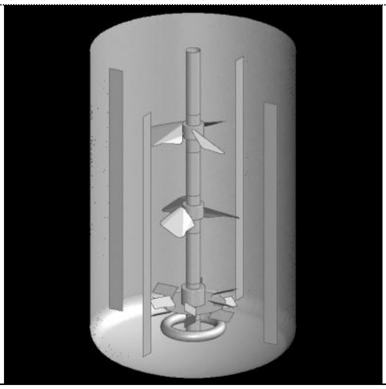 |

$^{\dagger}$Viscosity $\mu$ was measured by viscometry; the data were fitted to the non-Newtonian power-law model, where $\gamma$ is the shear rate ($s^{-1}$).

**Table S4. Results of the CFD analysis.**

| Scale (L) | Strain | Agitation (rpm) | $V_{tip}$† (m s$^{-1}$) | $P_{gV}$ (kW m$^{-3}$) | Velocity (m s$^{-1}$) | $k_La$ (h$^{-1}$) | Shear rate (s$^{-1}$) | Shear stress (Pa) |
|---|---|---|---|---|---|---|---|---|
| 200 | AGΔ-GAGΔ | 300 | 4.16 | 1.53 | 0.310 | 33.9 | 17.5 | 30.1 |
| 200 | | 400 | 5.55 | 3.33 | 0.490 | 51.8 | 27.7 | 40.4 |
| 200 | | 446 | 6.19 | 4.50 | 0.610 | 58.5 | 33.7 | 42.3 |
| 200 | AGΔ-GAGΔ-Δ*rolA* | 270 | 3.75 | 1.17 | 0.427 | 28.6 | 20.3 | 19.3 |
| 200 | | 340 | 4.72 | 2.10 | 0.512 | 36.0 | 25.3 | 24.1 |
| 200 | | 390 | 5.41 | 3.21 | 0.660 | 47.6 | 33.9 | 30.8 |
| 3000 | | 177 | 2.46 | 0.946 | 0.606 | 28.2 | 9.58 | 26.4 |
| 3000 | | 223 | 3.09 | 1.88 | 0.803 | 37.9 | 15.2 | 35.5 |
| 3000 | | 263 | 3.65 | 3.21 | 0.949 | 47.1 | 20.8 | 51.6 |

† $V_{tip}$: Impeller tip speed (m s$^{-1}$).
